# Stable Somatotopy in Chronic Low Back Pain: Is Cortical Map Reorganization a Myth?

**DOI:** 10.1101/2025.11.05.686896

**Authors:** M Dörig, DM Cole, A Guekos, P Stämpfli, P Schuetz, L Schibli, P Schweinhardt, ML Meier

## Abstract

Chronic low back pain (CLBP) has been linked to maladaptive cortical plasticity in sensorimotor regions, which may contribute to pain persistence through less distinct (“smudged”) somatotopic maps of afferent input from the back. However, empirical support for such functional reorganization processes, especially in the primary somatosensory cortex (S1), is limited. To delineate precise sensory representations of the back and test for altered somatotopic processing in CLBP, we utilized an MR-compatible pneumatic vibration device and applied frequency-specific vibrotactile stimuli (20 Hz and 80 Hz) at nine thoracolumbar paraspinal sites in 45 patients with CLBP and 41 healthy controls. Representational similarity analysis (RSA) was employed via a whole-brain searchlight method, and the neural representational patterns were contrasted against theoretical models including a segmental (based on anatomical proximity), simple (upper vs. lower back division) and a random model. In addition, a machine learning classifier was trained to predict afferent input from the upper vs lower back in healthy controls and tested in CLBP patients. Both groups displayed well-organized segmental representations spanning somatosensory, motor, and posterior parietal cortices. Posterior parietal regions exhibited the best model fits, followed by S1. No evidence for differences in representational patterns between groups was found. In CLBP patients, these patterns did not show associations with pain duration, severity, fear of movement or self-reported back perception. The classifier, trained on healthy controls, accurately predicted upper vs lower back afferent input in CLBP patients. These findings indicate preserved cortical maps of the back in CLBP, challenging the hypothesis of sensory cortical reorganization in this condition.

**Significance Statement:** Chronic low back pain (CLBP) is often linked to maladaptive reorganization of somatotopic maps in sensorimotor cortices, providing the mechanistic rationale for therapies like sensory discrimination training to “normalize” these maps and alleviate pain. Using frequency-specific vibrotactile stimulation of the back and multivariate fMRI analysis, we found well-organized, segmental cortical representations in both CLBP patients and healthy controls, with no evidence of group differences. Representational patterns showed no associations with clinical measures such as pain duration or intensity. These findings challenge the cortical reorganization hypothesis in CLBP and suggest sensory training benefits may involve alternative mechanisms.

## 1. Introduction

Chronic low back pain (CLBP) is a common and debilitating disorder and a major cause of disability worldwide (Thiese et al., 2014; Hartvigsen et al., 2018; James et al., 2018). In most cases of CLBP, a specific nociceptive source cannot be clearly identified (Maher et al., 2017). Beyond its potential musculoskeletal origins, CLBP is increasingly being recognized to involve neuroplastic changes at the supraspinal level, including alterations in cortical structure, function, and connectivity within sensorimotor and pain-related networks (Apkarian et al., 2011; Wand et al., 2011; Kong et al., 2013). Resting!zlstate fMRI analyses indicate that the primary somatosensory cortex (S1) representation of the back becomes hyperconnected with salience and default!zlmode networks, and that these connectivity patterns vary with pain intensity and catastrophizing in CLBP patients (Kim et al., 2019). Structural MRI studies report increased gray!zlmatter volume in S1 and the secondary somatosensory cortex (S2), as well as decreased white matter fractional anisotropy in S1, suggesting microstructural remodeling in CLBP (Kim et al., 2020; Medrano-Escalada et al., 2022). Moreover, CLBP is linked to altered excitability of sensorimotor cortices (Tsao et al., 2008; Jenkins et al., 2022). Beside these supraspinal alterations on structural and network levels, a somatotopic-specific functional cortical reorganization of the S1 back representation has been proposed to be linked to CLBP (Flor et al., 1997; Lloyd et al., 2008). Flor et al. (1997) provided the first evidence for S1 somatotopic reorganization in CLBP using magnetoencephalography (MEG). They reported a medial shift in the pain-evoked S1 response for the back representation, with larger shifts in patients with longer pain duration. Subsequent psychophysical work showed elevated two-point discrimination thresholds and distorted body outlines of the lower back in CLBP patients, interpreted as evidence of a cortical somatotopic shift or “blurring” in S1 (Moseley, 2008a; Adamczyk et al., 2018). These observations provided a mechanistic framework for the training of patients in refining their body image and tactile discrimination of the lower back, hypothesized to normalize cortical representations and reduce pain (Luomajoki and Moseley, 2011; Catley et al., 2014; Brumagne et al., 2019).

However, attempts to replicate the original findings of S1 reorganization in CLBP (Flor et al., 1997) have been limited and yielded mixed results. One study reported no somatotopic changes in S1 but found alterations in the secondary somatosensory cortex (S2) (Hotz-Boendermaker et al., 2016). Lloyd et al (2008) observed a medial S1 shift of the back representation during tactile stimulation in CLBP patients with high pain-related illness behavior (Waddell signs), suggesting an association between psychological factors and somatotopic S1 changes. A key limitation is that earlier CLBP neuroimaging research lacked fine-grained maps of the back’s somatotopic organization in S1 to assess potential spatial reorganization of paraspinal afferent input. In addition, most prior fMRI studies used univariate analyses of signal magnitude differences, which are relatively insensitive to subtle changes in representational structure (Popal et al., 2020). Advanced multivariate techniques, such as representational similarity analysis (RSA), offer greater sensitivity for probing cortical map structure (Kriegeskorte et al., 2008).

Recent neuroimaging studies have begun to address these limitations. Cole et al. (2022) introduced and validated an MR-compatible pneumatic vibration device (pneuVID) to deliver vibrotactile stimulation at different frequencies across multiple thoracolumbar levels, enabling the first mapping of the sensory cortical representation of paraspinal afferent input along the thoracolumbar axis. Using this device in combination with RSA, Guekos et al. (2023) demonstrated an orderly segmental cortical map in healthy individuals, such that adjacent paraspinal levels evoked more similar neural patterns than distant levels, reflecting a dermatomal-like organization.. Notably, this orderly map was evident not only in S1, but also in other sensorimotor regions (e.g., the superior parietal lobule (SPL) and primary motor cortex (M1)), suggesting a distributed network sharing the organizational structure of thoracolumbar somatosensory input. These previous studies established an important basis against which potential cortical somatotopic reorganization can be rigorously tested in patients with CLBP, which was the aim of the current study.

## 2. Materials and Methods

### 2.1 Experimental Design

The experiment consisted of a single fMRI session during which vibrotactile stimulation was delivered to the participants’ back. The protocol used a between!zlsubjects design. Ethical approval was granted by the ethics board of the Canton of Zurich, and all procedures conformed to the Declaration of Helsinki. Written informed consent was obtained from all participants prior to their inclusion in the study.

### 2.2 Participants

A total of 41 healthy adults (25 females) aged 21 to 39 years (mean age = 29.6 years, SD = 5.2 years) with a body mass index (BMI) below 30 kg/m² (mean BMI = 22.4 kg/m², SD = 2.6 kg/m²) and 45 patients with CLBP (28 females) aged 18 to 47 years (mean age = 30.5 years, SD = 8.3 years) with a BMI below 30 kg/m² (mean BMI = 22.8 kg/m², SD = 3.2 kg/m²) were included.

Inclusion criteria for patients with CLBP were a history of low back pain lasting more than three months, with no clinical indications of “red flags” such as infection, trauma, fractures, or inflammatory spondyloarthropathies. Exclusion criteria for healthy participants included any back pain within the past three months. Exclusion criteria common to both groups were other musculoskeletal pain within the past three months, any history of chronic pain (except CLBP in the patient group), spine, foot, or ankle surgery; psychiatric or neurological disorders; and excessive alcohol consumption (≥2 standard glasses/day for women, ≥4 standard glasses/day for men), use of intoxicants, analgesic intake within the past 24 hours, motor system impairments, and MRI contraindications.

Patients were asked to fill out several questionnaires: The German versions of the painDETECT (Freynhagen et al., 2006), State and Trait Anxiety Questionnaire (STAI) (Spielberger, 1971), Tampa Scale of Kinesiophobia (TSK) (Rusu et al., 2014), Oswestry Disability Index (OD) (Fairbank and Pynsent, 2000), and the Fremantle Back Awareness Questionnaire (FreBAQ) (Wand et al., 2016).

### 2.3 Pneumatic Vibration Device (pneuVID)

#### 2.3.1 pneuVID system

A custom-built pneumatic MR-compatible stimulation device (*pneuVID*) was used to deliver localized vibrotactile stimulation. PneuVID consists of manually engineered silicone actuators, which allows for safe operation during concurrent MRI acquisition without introducing imaging artifacts. Compressed air pulses at a constant pressure of 1.5 bar were transmitted through a valve box located at the foot of the MR scanner bed to achieve amplitudes between 0.5 to 1 mm. The stimulation was controlled by a Raspberry Pi-based module (Raspberry Pi Holdings plc, Cambridge, UK), connected to the valve box via a fiber-optic link to ensure electromagnetic compatibility with the MRI system. Further technical specifications, component design, and integration details have been previously described (Schibli et al., 2021; Cole et al., 2022).

#### 2.3.2. Stimulation Protocol

Prior to placement of the stimulation units, nine thoracolumbar vertebral levels (T3, T5, T7, T9, T11, L1, L3, L5, S1; T = thoracic, L = lumbar, S = sacral) were identified via palpation by an experienced physiotherapist. The vibrotactile units were positioned bilaterally to these locations along the erector spinae muscles, as well as to the muscle-tendon junction of the triceps surae muscle (mTS), using a sugar-based glue developed in-house and adhesive tape. Prior to brain imaging, a sensory testing procedure was undertaken to confirm that the vibrotactile stimuli were consistently perceived at each segmental level, with sensations localized centrally along the vertebral axis.

Stimulation was applied at two frequencies, 20 Hz and 80 Hz, based on their proposed generation of differential activation profiles of tactile and proprioceptive afferents: Lower frequencies are known to preferentially stimulate superficial mechanoreceptors such as Meissner corpuscles, while higher frequencies are more effective in activating more deeply located proprioceptors such as muscle spindles (Weerakkody et al., 2007; Avanzino et al., 2014a; Schellekens et al., 2021). However, this stimulation–response relationship is not strictly frequency-specific; for instance, Pacinian corpuscles, which respond to a broad frequency range (approximately 50–400 Hz) and have been shown to convey proprioceptive information via skin stretch (Talbot et al., 1968; LaMotte and Mountcastle, 1975; Weerakkody et al., 2007; Chung et al., 2013), may also be co-activated during 80 Hz stimulation (Avanzino et al., 2014a; Schellekens et al., 2021; Guekos et al., 2023).

During the experiment, each participant completed two functional runs, each containing 120 stimulation events (10 locations × 2 frequencies × 6 repetitions). Each stimulation event lasted for 5 seconds and was followed by a jittered interstimulus interval ranging from 4 to 7 seconds. The order of stimuli was pseudorandomized for each participant, ensuring no more than three consecutive events at the same location or frequency.

### 2.4 Neuroimaging Methods

#### 2.4.1 Image acquisition

Functional and structural imaging data were acquired using a 3 Tesla Philips Achieva MRI scanner (Philips, Best, The Netherlands) equipped with a 32-channel SENSE-compatible head coil. The stimulation protocol was split into two runs of approximately 22.5 minutes each. For each run, 750 T2*-weighted echo-planar images (EPIs) were collected, capturing blood-oxygen-level-dependent (BOLD) signals (TR = 1.8 s; TE = 34 ms; flip angle = 70°). The functional images consisted of 54 axial slices acquired in an interleaved ascending order, with a multiband factor of 3. In-plane spatial resolution was 1.72 × 1.72 mm², slice thickness was 2.0 mm with no inter-slice gap. Parallel imaging was applied using a sensitivity encoding (SENSE) factor of 1.4 and an EPI factor of 87. The scanner automatically discarded initial dummy volumes to allow for signal equilibration. Between the two functional runs, a high-resolution T1-weighted anatomical image was acquired using an MPRAGE sequence (TR = 6.6 ms; TE = 3.1 ms; flip angle = 9°; field of view: 230 × 226 × 274 mm; voxel size: 1.0 × 1.0 × 1.2 mm^3^; turbo field echo factor = 203). The complete imaging session lasted approximately 50 minutes per participant.

#### 2.4.2. Image Pre-processing

Following previous work (Cole et al., 2022), functional and anatomical MRI data were preprocessed using *fMRIPrep* (v20.2.0) (Esteban et al., 2019). T1-weighted images were corrected for intensity non-uniformity, skull-stripped using Advanced Normalisation Tools ANTs (v2.1.0) (Tustison et al., 2010), and segmented into gray matter, white matter, and cerebrospinal fluid using FSL FAST (Zhang et al., 2001). Brain surfaces were reconstructed with FreeSurfer (v6.0.1), and spatial normalization to MNI152NLin6Asym space was performed via nonlinear registration using ANTs. Motion correction of the functional data was implemented using the MCFLIRT tool (FSL) (Smith et al., 2004), and were co-registered to the T1w image using boundary-based registration (Greve and Fischl, 2009). These transformations were applied in a single step using ANTs (using Lanczos interpolation) to produce normalized, unsmoothed BOLD time series.

Regression based on confounders was performed on these preprocessed time series using a combination of tissue-based signals (mean, squared, and first derivatives from CSF and WM) and ICA-based components identified as motion-related artifacts using ICA-AROMA (Pruim et al., 2015), Aggressive denoising was applied using *fsl_regfilt*. No temporal filtering was applied. This procedure resulted in denoised data for each stimulation run, suitable for participant-level statistical analysis.

#### 2.4.2. First-Level Analysis

First-level analyses were performed using FSL FEAT (v6.00) with FILM pre-whitening enabled. The denoised fMRI time series were temporally high-pass filtered (100 s cutoff), spatially smoothed using a 4 mm FWHM Gaussian kernel, and analysed in standard space (2 mm isotropic voxels). For each run, the stimulation paradigm was modelled with 20 explanatory variables (EVs) corresponding to the 9 paraspinal stimulation locations and the triceps surae (mTS), as well as 2 stimulation frequencies (20 Hz and 80 Hz). The model used the precise onsets and durations (5 s) of each stimulation block. These were convolved with a canonical Gamma haemodynamic response function.

The general linear model included several contrasts of interest: (i) 80 Hz > 20 Hz stimulation across the nine paraspinal locations (excluding mTS); (ii) the inverse contrast; (iii) upper (T3–T11) > lower (L1–S1) stimulation sites; and (iv) the inverse. Additional contrasts were computed for each EV to isolate frequency-location-specific responses.

For each participant, statistical contrast images (contrast of parameter estimates; COPEs) were extracted for the nine paraspinal stimulation locations at both frequencies. These served as inputs for a subsequent representational similarity analysis (RSA), following the procedure described in Cole et al. (2022).

### 2.5. Representational Similarity Analysis (RSA)

RSA is a multivariate analysis technique used to investigate how patterns of neural activity represent different types of stimuli (Kriegeskorte et al., 2008). Unlike traditional univariate approaches, which average the signal across voxels and assess activation magnitude, RSA preserves the fine-grained multivoxel activity pattern. It analyzes the structure of neural representations by comparing activation patterns across conditions, offering insights into the geometric layout of stimulus representations in brain space (Dimsdale-Zucker and Ranganath, 2018; Popal et al., 2020).

The core element of RSA is the representational dissimilarity matrix (RDM), a symmetrical matrix that quantifies the dissimilarity between neural response patterns for all pairs of stimuli. Typically, the matrix contains zeros along its diagonal (indicating perfect similarity of a condition with itself), with increasing dissimilarity values representing more distinct activation patterns between conditions. The dissimilarity can be computed using several distance metrics, such as Pearson or Spearman correlation, Euclidean distance, or Mahalanobis distance.

#### 2.5.1. Software

RSA was implemented using Jupyter Notebooks (version 1.0.0, (Jupyter, 2015)) within the University of Zurich’s ScienceCloud supercomputing environment, running Neurodesk (v2024-04-04; Renton et al., 2024). The analysis was carried out with Python version 3.11.6 (Van Rossum and Drake, 2009) and the Python Representational Similarity Analysis toolbox rsatoolbox (version 0.0.4, Kriegeskorte et al., 2021).

The analysis also relied on several additional Python packages, including Nipype (version 1.8.6, Gorgolewski et al., 2011) interfacing FSL (FSL version 6.0.7.4, FMRIB Software Library, University of Oxford, UK), NiBabel (version 5.3.2, Brett et al., 2022) for loading and processing neuroimaging data, Nilearn (version 0.12.1, Abraham et al., 2014) for statistical analysis, machine learning and visualization, and scikit-learn (version 1.7.2, (Pedregosa et al., 2011) for machine learning analyses. Additional packages included siibra-python, (version 1.0.1a2, Dickscheid et al., 2025) to quantify the anatomical distribution in specific brain areas and NLTools (version 0.4.5, Chang et al., 2021) for various neuroimaging data manipulation. For data handling, manipulation and visualizations, the Pandas package (version 2.3.2, McKinney and Team, 2022), NumPy (version 1.26.4, Harris et al., 2020), and Matplotlib (version 3.8.4, Hunter, 2007) were used.

#### 2.5.2 Brain RDMs

For each participant, a 9 × 9 brain representational dissimilarity matrix (RDM) was generated for every contrast. Each RDM captures the dissimilarity between neural activation patterns across all pairs of experimental conditions, that is, between every pair of stimulated paraspinal locations. In this study, dissimilarity was quantified using the crossvalidated Mahalanobis distance (Crossnobis), which estimates the geometric distance between activation patterns while accounting for noise covariance and employing data partitioning for crossvalidation (Kriegeskorte et al., 2006; Walther et al., 2016). Crossnobis has been demonstrated to be the most reliable dissimilarity measure for multivoxel pattern analysis (Walther et al., 2016).

#### 2.5.3 Model RDMs

To interpret the organizational structure of neural activation patterns evoked by paraspinal vibrotactile stimulation, three theoretical model RDMs were constructed. These models reflect different hypotheses about how spatially organized sensory input from different back segments is represented in the brain.

##### 1. Simple Model

The simple model provides a categorical distinction between upper and lower back stimulation. Segments are divided into two regions: thoracic (Th3–Th11) and lumbosacral (L1–S1). It assumes that activation patterns within each region are indistinguishable (dissimilarity = 0), but entirely distinct across regions (dissimilarity = 1). This model tests whether the brain broadly separates sensory input from the upper and lower back, without encoding finer segment-level differences.

##### 2. Segmental Model

The segmental model encodes a graded representation of paraspinal sensory input based on the anatomical distance between stimulation sites. It assumes that activation patterns evoked by stimulation of neighboring spinal segments (e.g., Th3 and Th5) are spatially more similar than those from distant segments (e.g., Th3 and L5), in line with dermatomal organization (Downs and Laporte, 2011). Accordingly, dissimilarity values in the RDM increase stepwise by 0.1 with increasing segmental distance. An exception is the L5-S1 pair, which was set to a lower dissimilarity value (0.05) to reflect their closer physical proximity on the back. This model captures fine-grained spatial distinctions in the neural coding of sensory input.

##### 3. Random Model

The random model represents a null hypothesis of arbitrary brain responses, with no systematic structure related to the stimulated spinal segment. It serves as a control in model comparison analyses and contains no meaningful information about spatial organization. Dissimilarity values in this RDM are assigned randomly and do not reflect any anatomical or functional logic.

Together, these three model RDMs allow for a graded evaluation of how well the neural pattern data align with theoretical predictions, ranging from detailed spatial encoding (segmental model) to broad regional distinction (simple model).

#### 2.5.4. Searchlight approach

RSA can be combined with a searchlight approach, a method that systematically explores local patterns of activity across the brain (Kriegeskorte et al., 2006; Nili et al., 2014). A spherical “searchlight” of voxels is moved across pre-defined brain regions - such as the whole brain or a region of interest. At each position, RSA is performed on the local pattern within the searchlight, producing a similarity-based map of how well each area conforms to a theoretical model.

In this study, a whole-brain spherical searchlight analysis was conducted with a diameter of 5 voxels, resulting in a total of 155,815 searchlights. At each searchlight location, the neural RDM was compared to the three theoretical model RDMs using Kendall’s rank correlation coefficient (), a non-parametric and robust similarity metric particularly suited for models with tied ranks (Nili et al., 2014; Popal et al., 2020). The resulting correlation coefficients, one per searchlight, were Fisher-z transformed for statistical analysis (Walker, 2003).

Group-level inference was performed using FSL Randomise with 10,000 permutations to generate null distributions. Threshold-Free Cluster Enhancement (TFCE) was applied to identify significant clusters, with family-wise error (FWE) controlled at 5% (Smith and Nichols, 2009). This procedure was used for all second-level analyses. For within-group analyses, model fit comparisons (segmental vs. simple model and segmental vs. random model) were assessed using z-score difference maps followed by one-sample t-tests and TFCE correction. These analyses were conducted separately for 20 Hz and 80 Hz stimulation. Moreover, to assess potential relationships between the organizational structure of brain activity patterns and clinically relevant variables, regression analyses were performed within the CLBP group. These analyses examined z-score difference maps between the segmental and random models at both 80 Hz and 20 Hz stimulation frequencies. Regressors included pain duration (in days), FreBAQ scores, TSK scores, current pain immediately before the MRI session and maximal and average pain over the past four weeks (assessed using an 11-point Numerical Rating Scale, NRS). Separate contrasts were specified to test for positive and negative associations with each behavioral regressor.

For between-group comparisons, potential differences between healthy controls and patients with CLBP were assessed using unpaired two-sample t-tests. This analysis was performed on both individual model fits and model comparison maps (z-score difference maps), separately for each stimulation frequency (20 Hz and 80 Hz).

### 2.6. Machine Learning Analysis

A linear support vector machine (SVM) classifier was trained to distinguish between upper (thoracic) and lower (lumbosacral) back stimulation, using COPEs from all stimulation sites within each region at both frequencies (20 and 80 Hz). The classifier was trained on the healthy control group and tested on the CLBP group to compare classification accuracy between groups.

Analyses were carried out within a region-of-interest (ROI) mask encompassing the sensorimotor cortex, primarily including Brodmann areas 1–5 (primary somatosensory cortex, primary motor cortex, and superior parietal lobule). The ROI mask was confined to the brain by intersecting it with a standard MNI brain mask (as provided in Nilearn), ensuring that only voxels within the brain were included. Voxel intensities within the mask were standardized before classification.

Within the healthy controls, model selection and performance estimation were performed using nested cross-validation, with a 3-fold inner loop to optimize the penalty parameter *C* of the linear SVM and a 5-fold outer loop to estimate generalization accuracy. Feature selection was performed using ANOVA F-scores to identify the most discriminative voxels within the ROI. Accuracy was chosen as the scoring metric, as both classes (thoracic vs. lumbosacral stimulation) contained equal numbers of samples. Classification performance was quantified using accuracy, complemented by confusion matrices and classification reports, and statistical significance was assessed using 5000-label permutation testing.

Finally, we derived weight maps from the trained linear SVM to visualize which brain regions contributed most strongly to the discrimination of thoracic versus lumbosacral stimulation.

SVM coefficients were transformed back into brain space and visualized on the MNI152 template, thresholded at the 99th percentile of absolute weight values. For anatomical localization and reporting, discriminative voxels were characterized using the Harvard-Oxford Cortical Structural Atlas (Desikan et al., 2006), and, for cytoarchitectonic labeling, the probabilistic JuBrain atlas as implemented in the SPM Anatomy Toolbox (Eickhoff et al., 2005; Zilles and Amunts, 2010).

### Code/Software

Jupyter notebooks including the code for RSA analysis and machine learning are available on GitHub (https://github.com/ISR-lab/RSA).

## 3. Results

### 3.1 Baseline characteristics of study sample

Mean age was 29.6 years in healthy controls and 30.5 years in CLBP patients. There were no significant differences between healthy controls and CLBP patients regarding age or BMI (p’s > 0.5) or the proportion of female participants (62.2% in healthy controls vs. 61.0% in patients; χ² = 0.0141, p = 0.905). Self-perception of the back, as measured by the FreBAQ, was significantly different between groups (p < 0.001), suggesting body image distortion in patients with CLBP (Wand et al., 2016). For a complete overview of the cohort characteristics, refer to Table 1.

**Table 1.** Baseline characteristics.

|  | *Healthy Controls* | *CLBP Patients* | *Test statistics* |
| --- | --- | --- | --- |
| ***n*** | 41 | 45 |  |
| ***Age (years)*** | 29.60 [21.00, 39.00] | 30.5 [18.00, 47.00] | $t = 0.610$, $p = 0.543$ (a)<br>Mann-Whitney $U = 911$,<br>$p = 0.924$ |
| ***Gender (n female (%))*** | 25 (62.22) | 28 (60.98) | $\chi^2 = 0.0141$, $p = 0.905$ |
| ***Weight (kg)*** | 66.30 [44.90, 92.30] | 69.00 [44.80, 99.70] | $t = 0.994$, $p = 0.323$ |
| ***Height (cm)*** | 172.00 [154.00, 187.00] | 173.00 [153.00, 195.00] | $t = 0.847$, $p = 0.400$ |
| ***BMI (kg/m<sup>2</sup>)*** | 22.40 [18.80, 29.10] | 22.8 [17.20, 30.80] | $t = 0.650$, $p = 0.518$ |
| ***Fremantle Back Awareness Questionnaire (FreBAQ)*** | 1.71 [0.00, 11.00] | 7.22 [0.00, 27.00] | $t = 5.863$, $p < 0.001^*$ (a)<br>Mann-Whitney $U = 233$,<br>$p < 0.001^*$ |
| ***Tampa scale of kinesiophobia*** | 31.50 [19.00, 44.00] | 32.40 [22.00, 49.00] | $t = 0.694$, $p = 0.489$ |
| ***Oswestry disability index*** | NA | 28.90 [18.00, 52.00] |  |
| **painDetect questionnaire** | NA | 14.30 [4.00, 25.00] |  |
| **Momentary pain at MRI session** | NA | 2.36 [0.00, 6.00] |  |
| **Average pain during the past 4 weeks** | NA | 3.78 [1.00, 7.00] |  |
| **Maximum pain during the past 4 weeks** | NA | 6.31 [1.00, 10.00] |  |
| **Pain duration (days, self-reported)** | NA | 2786 [150, 7849] |  |
| *N(%, mean [min, max].*<br>a) *Independent Samples T-Test: Levene's test is significant ($p < 0.05$), suggesting a violation of the assumption of equal variances* |  |  |  |

### 3.2 RSA results

#### 3.2.1 Within-group comparisons

##### Segmental > random models at 80 Hz

To assess whether the brain activity patterns reflect true thoracolumbar anatomical spacing rather than random variation, the segmental model was compared to the random model within each group.

In healthy controls, SPL areas exhibited the greatest overlap with voxels showing significantly stronger model fits to the segmental vs random model, led by area 5L (32.16%) and followed by 7pc (23.39%), 5ci (17.78%), 5m (13.43%), and 7a (7.78%). Within S1, overlap was strongest in area 2 (17.64%), followed by 3b (11.45%) and 3a (6.86%), with minimal overlap in area 1 (3.76%). M1 areas showed less overlap: 4p (2.66%) and 4a (2.09%), while premotor area 6d1 showed low overlap (0.04%).

Patients with CLBP demonstrated a similar regional pattern compared to healthy controls, with SPL again showing the greatest overlap: 5L (21.03%), 5m (12.84%), 5ci (12.34%), 7pc (9.72%), and 7a (3.55%). Within S1, overlap was observed in areas 2 (10.28%), 3b (7.67%), and 3a (3.63%), with minimal overlap in area 1 (2.29%). M1 areas showed low overlap as well: 4p (3.08%) and 4a (1.14%), while premotor area 6d1 showed no overlap (0.0%). The spatial distribution of significant effects in both groups and their overlap is shown in Figure 3.

**Figure 1.**
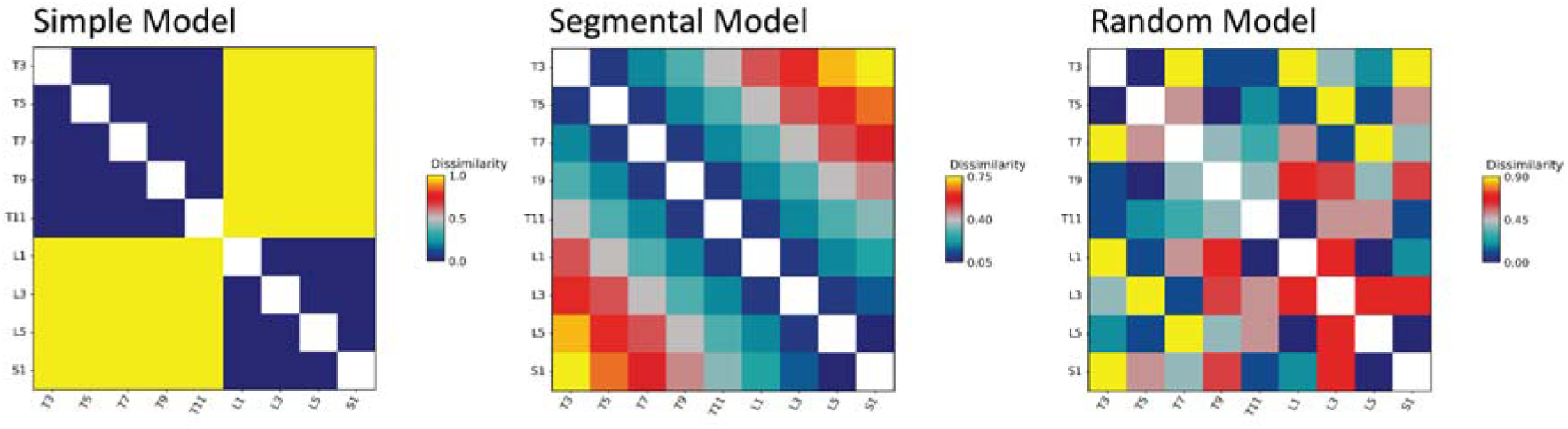
Three model RDMs were used in the RSA analysis. Color matrices illustrate dissimilarity patterns between spinal segments T3-S1 for the simple, segmental, and random models. Blue indicates low dissimilarity (high similarity) while red/yellow indicates high dissimilarity. The simple model shows binary thoracic-lumbar structure, the segmental model displays graded patterns based on segment proximity, and the random model lacks systematic organization.

**Figure 2.**
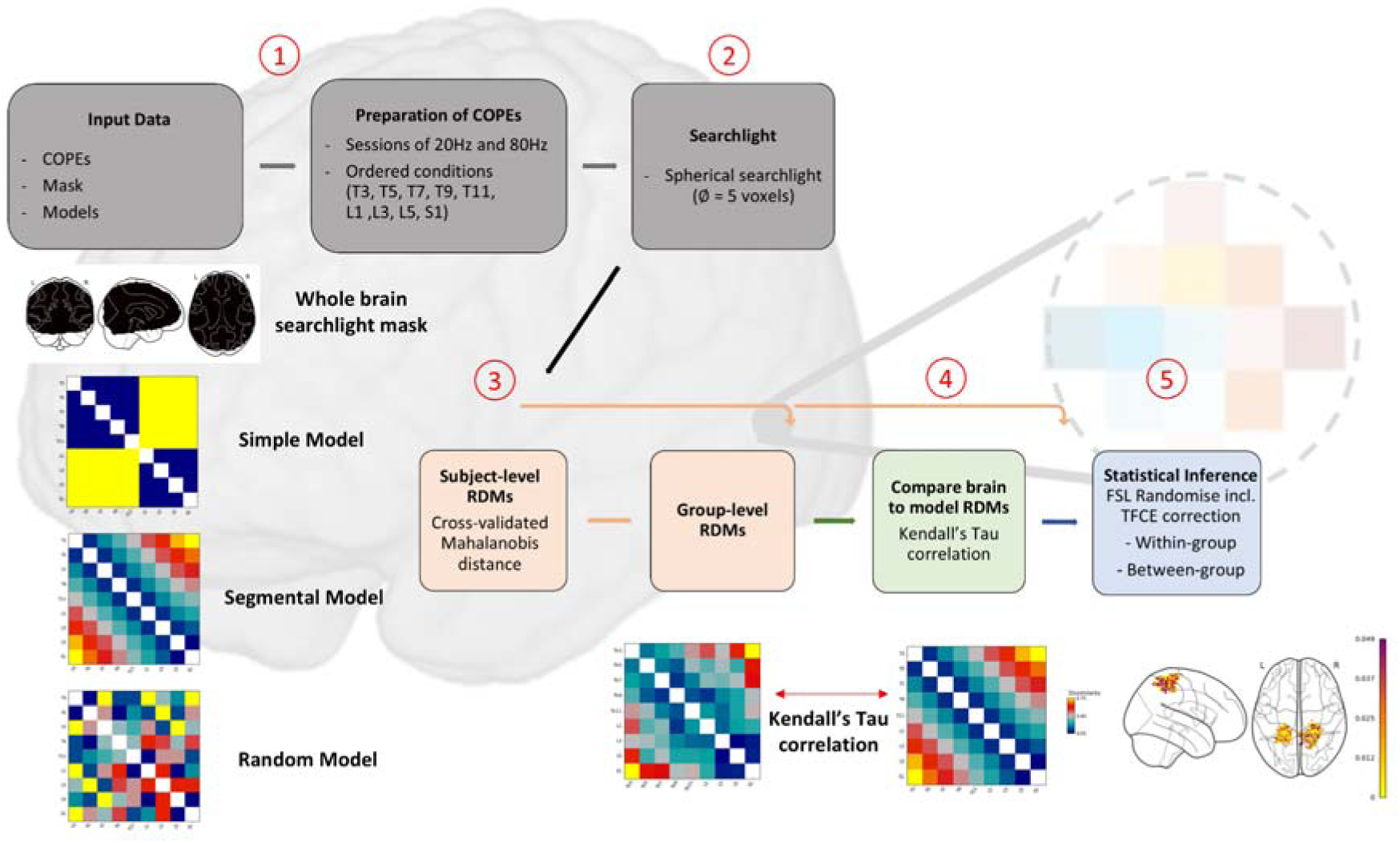
Overview of the analysis pipeline for the searchlight RSA. The workflow included: (1) data import and preparation of statistical contrast images, (2) execution of the searchlight procedure, (3) computation of individual and group-level brain representational dissimilarity matrices (RDMs) using Crossnobis distance, (4) comparison of brain RDMs to model RDMs using Kendall’s Tau correlation, and (5) statistical inference using FSL Randomise with Threshold-Free Cluster Enhancement (TFCE).

**Figure 3.**
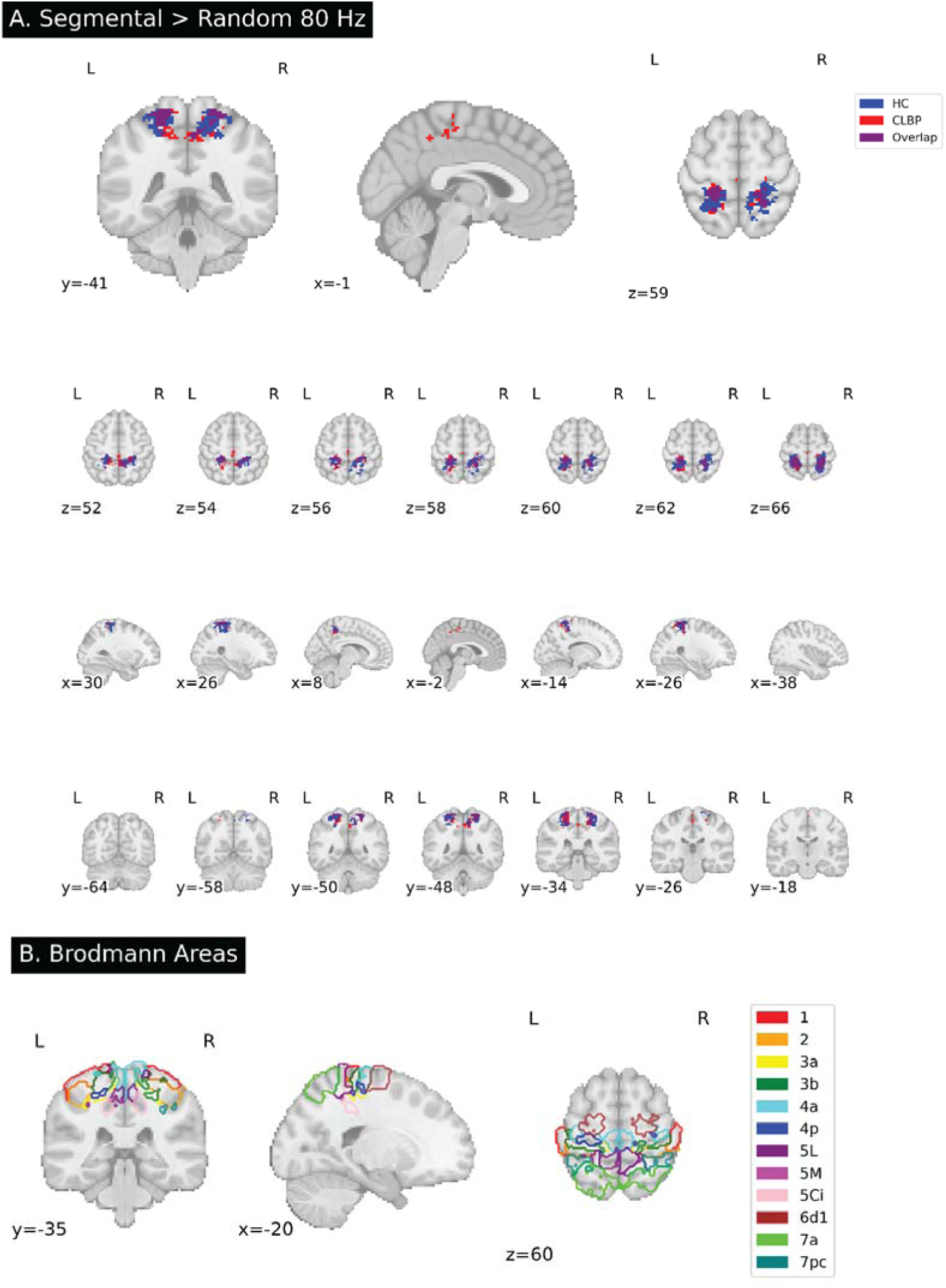
**A.** Segmental versus random model comparison at 80 Hz (within-group analyses). Significant clusters from one-sample t-tests on z-score difference maps (segmental > random), TFCE-corrected at p < 0.05 FWE. Blue: Healthy controls (N=41); Red: CLBP patients (N=41); Purple: overlap. Upper: orthogonal view. Lower: axial mosaic, both in MNI space. **B.** Corresponding Brodmann Areas (BA 1, 2, 3a, 3b, 4a, 4p, 5L, 5M, 5Ci, 6d1, 7a, 7pc) illustrating

##### Segmental > simple models at 80 Hz

This comparison tested for brain activity patterns encoding fine-grained segmental information beyond a simple upper vs. lower back distinction.

In healthy controls, most voxels showing significantly stronger model fits to the segmental vs simple model overlapped primarily with SPL areas, with area 5L demonstrating the greatest overlap (21.15%), followed by area 2 (12.81%) and 7pc (9.49%). Moderate overlap was observed in areas 3b (7.55%), 3a (7.27%), 5m (4.8%), and 7a (3.15%), while smaller differences were found in areas 5ci (2.93%), 1 (2.74%), 4p (2.38%), and 4a (1.33%). The premotor area 6d1 showed minimal overlap (0.04%).

Patients with CLBP showed reduced overlap across most regions in this comparison. Overlaps were observed in area 7pc (6.1%), followed by area 2 (2.45%) and 5l (1.2%). Most other regions showed minimal overlap: areas 1 (0.7%), 3b (0.24%), and 7a (0.12%) showed minimal overlap, while areas 3a, 4a, 4p, 5m, 5ci, and 6d1 covered no voxels that showed a significantly stronger model fit to the segmental vs simple model. The spatial distribution of significant effects in both groups and their overlap is shown in Figure 4.

**Figure 4.**
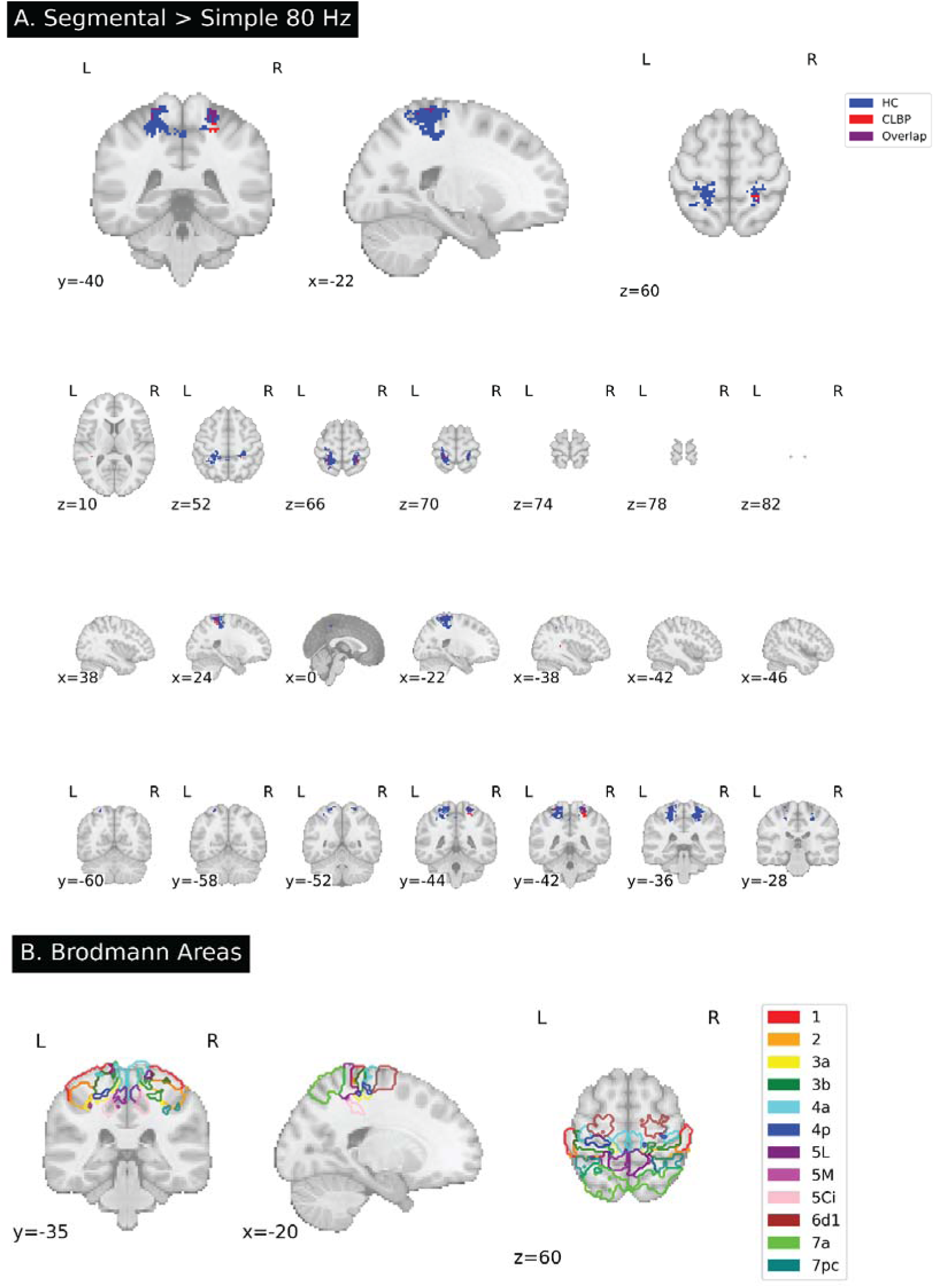
**A.** Segmental versus simple model comparison at 80 Hz (within-group analyses). Significant clusters from one-sample t-tests on z-score difference maps (segmental > random), TFCE-corrected at p < 0.05 FWE. Blue: Healthy controls (N=41); Red: CLBP patients (N=41); Purple: overlap. Upper: orthogonal view. Lower: axial mosaic, both in MNI space. **B.** Corresponding Brodmann Areas (BA 1, 2, 3a, 3b, 4a, 4p, 5L, 5M, 5Ci, 6d1, 7a, 7pc) illustrating

##### Model comparisons at 20 Hz

In healthy controls, the comparison of the segmental > random model yielded no significant voxels. Few voxels showed a significant model fit to the segmental model vs simple model. These were primarily located in the SPL (including area 7pc (5.65%), 5L (2.31%)), and area 2 (1.23%). In patients with CLBP, no significant voxels were detected for either comparison, segmentalL > random or segmentalL> simple.

#### 3.2.2. Regression analysis of the cLBP group

No significant positive or negative associations were found between the behavioral regressors (pain duration, FreBAQ scores, current pain, maximal and average pain over the past four weeks and TSK score) and the z-score difference maps at either 80 Hz or 20 Hz stimulation frequencies (p’s > 0.5).

#### 3.2.2. Between-group comparisons

No significant differences in representational structure were observed between patients with CLBP and healthy controls for any of the models tested (segmental, simple, or model contrasts) at either 20 Hz and 80 Hz (p’s > 0.05).

### 3.3 Machine learning results

Within the healthy control group, nested cross-validation yielded a mean accuracy of 0.73 (SD = 0.12) for discriminating upper vs lower back stimulation, with the best model obtained at a penalty parameter C=0.01. When applied to the patient group, the classifier achieved an accuracy of 0.78. Precision, recall, and F1-scores were comparable across the two classes (thoracic vs. lumbosacral), with macro-averaged values of 0.79 (precision), 0.78 (recall), and 0.78 (F1-score). The confusion matrix indicated higher sensitivity for thoracic stimulation (recall = 0.87) compared to lumbosacral stimulation (recall = 0.69). A permutation test (n = 5000) confirmed that the classification accuracy in the patient group was significantly above chance (p < 0.001).

Weight maps revealed the spatial distribution of discriminative voxels within the predefined sensorimotor ROI mask. Spatial characterization showed that most voxels were localized within S1, predominantly spanning Brodmann areas 3a and 3b, and the SPL, with minor overlap into area 4p (posterior primary motor cortex) (Figure 5).

**Figure 5.**
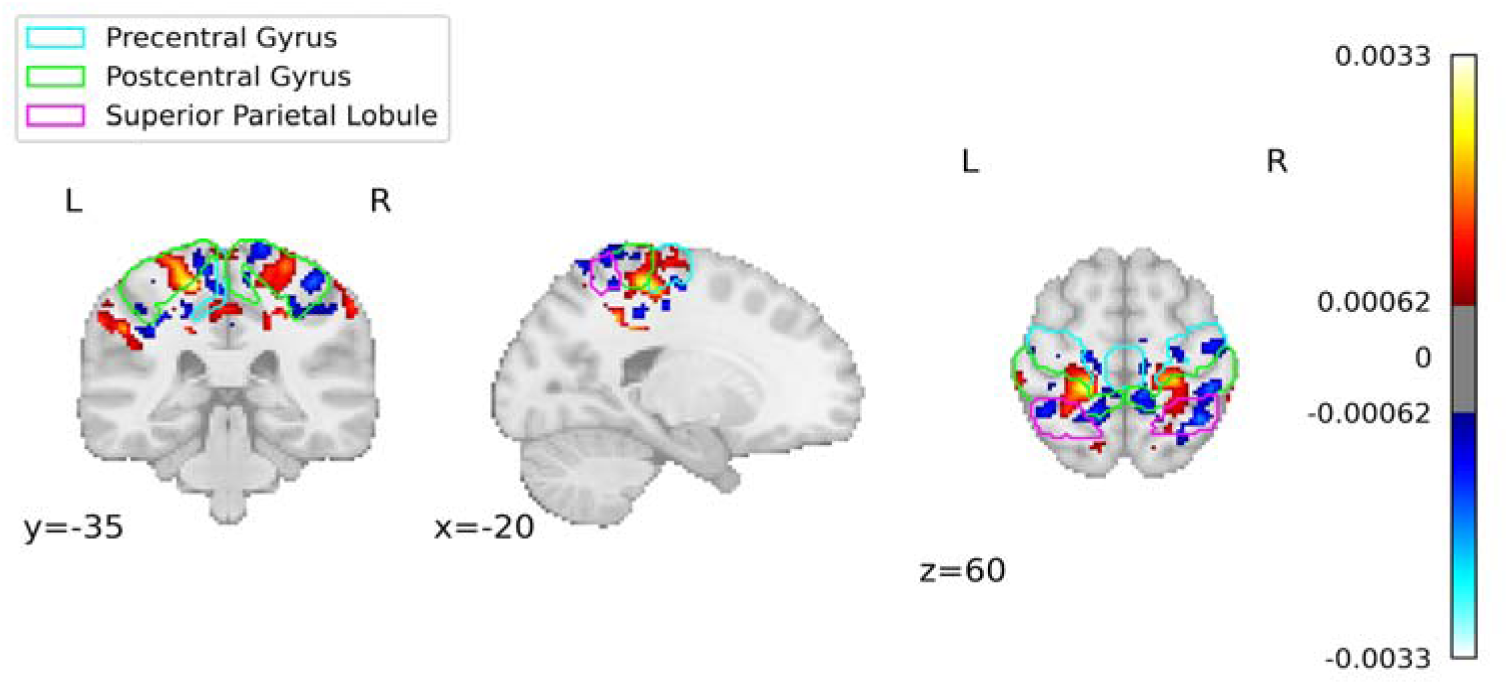
SVM weight map for upper vs lower back discrimination. Orthogonal slices showing classifier weights thresholded at the 99th percentile, projected onto the MNI152 template. Warm/cool colors indicate weights associated with thoracic/lumbosacral stimulation, respectively. Contours highlight the postcentral gyrus (magenta), precentral gyrus (lime), and superior parietal lobule (cyan).

Table 2 lists the 10 voxels with the highest absolute SVM weights, reporting the SVM weight, class support (thoracic or lumbosacral), MNI coordinates, and anatomical location using the JuBrain (SPM Anatomy Toolbox (Eickhoff et al., 2005)) probabilistic cytoarchitectonic atlas (Zilles and Amunts, 2010). The listed percentages indicate the probability that each voxel belongs to a given cortical area. The most contributing voxels were localized within left S1, predominantly spanning Brodmann areas 3a and 3b, with minor overlap into area 4p (posterior primary motor cortex).

**Table 2.** Ten voxels with the highest absolute SVM weights discriminating thoracic from lumbosacral stimulation. Values indicate SVM weight, class support, MNI coordinates, and probabilistic cytoarchitectonic labels from the JuBrain atlas. Percentages denote the probability of each voxel belonging to a given cortical area; *unclassifiable* voxels did not overlap with any labeled region.

| Rank | Weight | Class support | MNI Coordinates (xyz) | Region |
| --- | --- | --- | --- | --- |
| 1 | 0.003347 | thoracal (1) | -18.0 -34.0 58.0 | 21.2% Area 3a<br>21.2% Area 3b |
| 2 | 0.003274 | thoracal (1) | -16.0 -34.0 58.0 | 34.5% Area 3a<br>4.9% Area 3b |
| 3 | 0.003157 | thoracal (1) | -18.0 -32.0 58.0 | 40% Area 3a<br>3.2% Area 3b |
| 4 | 0.002975 | thoracal (1) | -18.0 -36.0 58.0 | 17.8% Area 3b<br>6.5% Area 3a |
| 5 | 0.002964 | thoracal (1) | -18.0 -34.0 60.0 | 27.6% Area 3b<br>11% Area 3a<br>3.4% Area 4p |
| 6 | 0.002930 | thoracal (1) | -18.0 -38.0 34.0 | unclassifiable |
| 7 | 0.002913 | thoracal (1) | -20.0 -32.0 58.0 | 33.4% Area 3a |
|  |  |  |  | 7.8% Area 4p<br>6.6% Area 3b |
| 8 | 0.002895 | thoracal (1) | -16.0 -36.0 58.0 | 16.6% Area 3a<br>10.0% Area 3b<br>1.1% Area 4p |
| 10 | 0.002883 | thoracal (1) | -20.0 -34.0 58.0 | 30% Area 3b<br>15.5% Area 3a<br>3.1% Area 4p |

## 4. Discussion

The present study employed thoracolumbar vibrotactile stimulation and leveraged multivariate pattern recognition methods to test for potential differences in fine-grained somatotopic cortical organization of paraspinal afferent input in patients with CLBP compared to healthy controls. Contrary to the longstanding assumption of maladaptive cortical map reorganization of afferent input from the back in S1 in CLBP (Flor et al., 1997), we found no evidence of somatotopic reorganization of the back in S1. As demonstrated with RSA, the fine-grained segmentally organized representational structure of paraspinal afferent input in CLBP was largely comparable to healthy controls. Importantly, within the CLBP group, the RSA patterns did not show associations with pain duration, pain intensity, fear of movement or self-reports of back perception suggesting resilient somatotopic maps of paraspinal afferent input. Moreover, the SVM classifier, successfully trained to predict upper vs lower back stimulation in healthy controls, generalized well to patients with CLBP, further supporting that cortical activity patterns related to thoracolumbar afferent input are preserved in CLBP. These findings do not only challenge the cortical map reorganization theory in CLBP but also raise doubts about the mechanistic rationale of interventions such as sensory discrimination training aiming at normalizing cortical organization to reduce pain (Moseley, 2008b; Catley et al., 2014; Kälin et al., 2016).

Beyond S1, segmentally well-organized representational patterns of thoracolumbar afferent input were also observed in both groups within the SPL and M1, indicating that spatial encoding of paraspinal afferent input extends beyond early somatosensory areas into higher-order multisensory networks and motor areas. The SPL is a key node of the dorsal posterior parietal cortex (PPC) known for integrating tactile and proprioceptive information into coherent body representations (Meier et al., 2019; Zeharia et al., 2019; Klautke et al., 2023). The SPL (in particular area 5) is closely interconnected with M1 through corticocortical projections (Beloozerova et al., 2022). These pathways enable proprioceptive and tactile information processed in SPL to modulate motor output and postural control (Naito et al., 2016; Mirdamadi et al., 2025). In these regions, the segmentally organized representations were more pronounced at 80 Hz stimulation in both groups, likely due to joint contributions of deeper (muscle spindles) and superficial proprioceptive (Pacinian corpuscles) and tactile (Meissner corpuscles) afferents (Avanzino et al., 2014b; Schellekens et al., 2021).

The current results align with emerging evidence questioning cortical map reorganization in chronic pain syndromes. An S1 mapping study in patients with unilateral complex regional pain syndrome (CRPS) reported that the cortical area, location, and geometry of the affected hand map were largely comparable to the unaffected hand and to healthy controls, with no link between somatotopic map measures and pain or disability (Mancini et al., 2019). The authors concluded that if reorganization occurs, it is not directly related to clinical severity, challenging the rationale for therapies in CRPS aimed at “restoring” S1 maps. Complementing these findings, recent work demonstrated that S1 maps remain remarkably stable even following major afferent loss. A longitudinal study tracked three individuals before and up to five years after arm amputation and found stable hand and lip representations in S1; pre–post comparisons showed no large-scale S1 reorganization (Schone et al., 2025). Together with the current findings, these data suggest that the somatotopic organization of the adult S1 is robust and that symptoms (e.g., pain, altered body perception) do not necessarily have to be accompanied by a disturbance of the topographic map in S1.

The current findings of stable somatotopic maps have important implications for interventions that target presumed cortical “smudging,” such as tactile or sensory discrimination training (Catley et al., 2014; Kälin et al., 2016). These therapies were originally developed under the assumption that chronic pain is partly maintained by degraded or overlapping S1 representations of afferent input from the back, which could be restored or normalized through repeated tactile stimulation training and spatial discrimination tasks. However, if the somatotopic maps remain functionally stable, as suggested by the present results, potential therapeutic benefits of sensory discrimination training (Kälin et al., 2016) likely arise from other mechanisms than direct reorganization of S1. Hence, rather than aiming to “restore” a disrupted map, future mechanistic investigations might focus on alternative cortical effects of sensory discrimination training such as changes in network connectivity, e.g. between S1 and SPL.

### Limitations

A methodological limitation of the present work is that cortical representations were probed using non-noxious vibrotactile stimuli rather than nociceptive input. Flor et al. (1997) originally employed brief nociceptive intradermal electrical stimuli to the back, based on the assumption that cortical reorganization in chronic pain is primarily driven by nociceptive input. Therefore, we cannot exclude the possibility that nociceptive maps of thoracolumbar input might show different organizational patterns. However, previous work demonstrated that nociceptive and tactile representations in S1 are largely co-localized. Namely, painful laser stimulation of individual fingers produced maps that were highly aligned with those evoked by non-painful air-puff stimulation, suggesting comparable cortical representations between mechanoreceptive and nociceptive signals (Mancini et al., 2012). Furthermore, CLBP is a heterogeneous condition which can be maintained by nociceptive, neuropathic, and nociplastic pain mechanism components (Shraim et al., 2022). Recent evidence suggests that a potential somatotopic reorganization may depend on the predominant pain mechanism. For example, S1 map reorganization seems to occur in patients with painful trigeminal neuropathy, but not in patients with a (non-neuropathic) painful temporomandibular disorder (Gustin et al., 2012) where nociplastic features often dominate (Harper et al., 2016). Furthermore, CLBP patients with predominant nociplastic features displayed less consistent changes in the M1 maps of back muscles than patients with both nociceptive and nociplastic features who showed greater overlap of back muscle representations (Elgueta-Cancino et al., 2021). Thus, heterogeneity of pain mechanisms in our study sample (mean PainDETECT score of 14.3 is suggesting unclear pain mechanisms [nociceptive vs neuropathic] (Packham et al., 2017)) could have diluted subgroup-specific effects. Future work should incorporate study designs suitable to distinguish between the different pain mechanisms (Shraim et al., 2021, 2022).

To conclude, using high-resolution vibrotactile mapping and advanced multivariate analyses, we found no evidence of cortical “smudging” or spatial disarray of thoracolumbar afferent input in CLBP, despite long-term pain and distorted body image of the back. These findings challenge the prevailing assumption that CLBP involves degraded S1 maps. From a clinical perspective, they call for reinterpretation of sensory-based interventions in CLBP. Future work should integrate longitudinal designs including pain mechanism identification and focus on other mechanisms of sensory-based interventions in CLBP such as changes in network connectivity within sensorimotor regions.

## Acknowledgments

This work was funded by the Swiss National Science Foundation through a grant to MM (grant number 320030_185123) and by the EMDO foundation (Switzerland). The authors wish to thank Magdalena Suter for helping with data collection and recruitment of participants.

